# Revisiting soil fungal biomarkers and conversion factors: Interspecific variability in phospholipid fatty acids, ergosterol and rDNA copy numbers

**DOI:** 10.1101/2024.01.04.574151

**Authors:** T Camenzind, H Haslwimmer, MC Rillig, L Ruess, DR Finn, CC Tebbe, S Hempel, S Marhan

**Affiliations:** Institute of Biology, Freie Universität Berlin, Altensteinstr. 6, 14195 Berlin, Germany; Berlin-Brandenburg Institute of Advanced Biodiversity Research (BBIB), Berlin, Germany; Institute of Soil Science and Land Evaluation, Soil Biology Department, University of Hohenheim, Stuttgart, Germany; Humboldt-Universität zu Berlin, Institute of Biology, Ecology group, Philippstraße 13, 10115 Berlin, Germany; Thünen Institut für Biodiversität, 38116 Braunschweig, Germany

## Abstract

The abundances of fungi and bacteria in soil are used as simple predictors for carbon dynamics, and represent widely available microbial traits. Soil biomarkers serve as quantitative estimates of these microbial groups, though not quantifying microbial biomass per se. The accurate conversion to microbial carbon pools, and an understanding of its comparability among soils is therefore needed. We refined conversion factors for classical fungal biomarkers, and evaluated the application of quantitative PCR (qPCR, rDNA copies) as a biomarker for soil fungi. Based on biomarker contents in pure fungal cultures of 30 isolates, combined with available references, we propose average conversion factors of 95.3 g fungal C g^−1^ ergosterol, 32.0 mg fungal C µmol^−1^ PLFA 18:2ω6,9 and 0.264 pg fungal C ITS1 DNA copy^−1^. As expected, interspecific variability was most pronounced in rDNA copies, though qPCR results showed the least phylogenetic bias. A modeling approach based on exemplary agricultural soils further supported the hypothesis that high diversity in soil buffers against biomarker variability, whereas also phylogenetic biases impact the accuracy of comparisons in biomarker estimates. Our analyses suggest that qPCR results cover the fungal community in soil best, though with a variability only partly offset in highly diverse soils. PLFA 18:2ω6,9 and ergosterol represent accurate biomarkers to quantify Ascomycota and Basidiomycota. To conclude, the ecological interpretation and coverage of biomarker data prior to their application in global models is important, where the combination of different biomarkers may be most insightful.

## Introduction

Soils fulfill crucial ecosystem functions like carbon (C) storage, plant nutrition and element cycling (Parikh and James, 2012). These functions are driven by a community of microorganisms, quantitatively typically dominated by bacteria and fungi, which also constitute a relevant component of C and nutrient cycles (Liang et al., 2019). However, the nature and complexity of soil makes it challenging to quantify microbial biomass. Direct microscopic counts are not only labor- and time-consuming, but also prone to error due to extraction inefficiencies, staining biases and subjectivity (Stahl et al., 1995; Joergensen and Wichern, 2008). Instead, biomarkers reflecting microbial biomass offer easy quantification and are used preferentially in many soil analyses. Phospholipid fatty acids (PLFAs) represent such standard biomarkers for soil microorganisms. Different organism groups show specific PLFA patterns – the major biomolecules in cell membranes (Frostegård and Bååth, 1996). The PLFA 18:2ω6,9 is used as a standard biomarker for saprotrophic and ectomycorrhizal fungi, while the PLFA 16:1ω5 is the common marker for arbuscular mycorrhiza fungi, and cyclic and branched chain PLFAs are frequent in different bacterial groups (Ruess and Chamberlain, 2010; Ngosong et al., 2012; Willers et al., 2015) As a consequence, the PLFA marker composition in soil provides insights into the relative abundance of these microbial groups.

Such group level resolution appears relatively coarse, but fungal:bacterial ratios represent important ecological indicators. Their ratios are discussed as a promising microbial trait to be included in biogeochemical models (He et al., 2021), also because of widely accessible datasets (Bar-On et al., 2018; Crowther et al., 2019; Yu et al., 2022). The relative abundances of fungi and bacteria correlate to soil parameters relevant for C cycling dynamics: high fungal:bacterial ratios have been shown to positively relate to C:N ratios, soil organic carbon stocks or litter decomposition rates (Strickland and Rousk, 2010; Malik et al., 2016; Yu et al., 2022). This pattern is driven by the special physiology of the mycelial growth form in fungi and their biochemical composition (e.g., C-rich cell walls and high C:N ratios; Mouginot et al., 2014; Moore et al., 2021). Fungal growth physiology together with their biochemical composition leads to high C- and nitrogen-use efficiencies (Camenzind et al., 2021), and likely also to the high contribution of fungal necromass to soil organic carbon stocks (Liang et al., 2019; Camenzind et al., 2023). This important role of fungi in soil C cycling demands more precise understanding for predicting their abundance based on biomarker-analyses.

Soil fungal C contents must be indirectly deduced from biomarker abundances via conversion factors. Biomarkers only represent a proxy for microbial biomass: in the first instance biomarkers are no more than quantitative values of this biomolecule in soil, with its primary origin from the specific organism group. Within the organisms, these biomolecules are only a fraction of the total biomass, and conversion of these proportional values to actual biomass is needed to correctly calculate C pools of respective groups. Such conversion factors are derived from relative biomolecule fractions in biomass (Joergensen and Wichern, 2008). However, especially in the case of the fungal PLFA marker linoleic acid (18:2ω6,9, hereafter referred to as fungal biomarker) limited data are available. Its general specificity for fungi has been demonstrated repeatedly: Federle (1986) was the first one to show that fungal mycelia are enriched in this specific PLFA (using 47 fungal species): the 18:2ω6,9 marker constituted on average 43% of total PLFAs (Frostegård and Bååth, 1996). Later, studies with further fungal isolates confirmed the specificity of this biomarker in fungi (Zelles, 1997; Klamer and Bååth, 2004; Taube et al., 2019). Other fatty acids frequently observed in fungal tissues are a(alpha)-linolenic acid (18:3ω3,6,9) and g(gamma)-linolenic acid (18:3ω6,9,12) (Weete, 1980; Vestal and White, 1989; Van der Westhuizen et al., 1994). However, these fatty acids are less specific and also found in higher plants (Ruess and Chamberlain, 2010).

Still, quantitative data of PLFA biomarker contents in fungal biomass are rare. Klamer and Bååth (2004) provided a first estimate for a conversion factor based on 12 fungal species isolated from compost: a value of 85 mg fungal C µmol^−1^ 18:2ω6,9 PLFA (as recalculated by Joergensen and Wichern, 2008). Baldrian et al. (2013) analyzed Basidiomycete mushrooms in forests, reporting a conversion factor of 12 mg fungal C µmol^−1^ 18:2ω6,9 PLFA based on 11 species. These data highlight the challenge to come up with one conversion factor for this biomarker: not only do studies diverge, also within studies large variation exists - the coefficient of variation (CV) among fungal isolates tested by Klamer and Bååth (2004) amounted to 74%. In addition, PLFA composition varies phylogenetically - especially Mucoromycota and Mortierellomycota show a specific PLFA composition (Zelles, 1997; Klamer and Bååth, 2004; Taube et al., 2019). The lipid pattern of fungi is variable even in systematically related fungi, *i.e*., taxonomic or phylogenetic relationships are not necessarily reflected in fatty acid profiles (Ruess et al., 2002). Consequently, fungal community shifts – as a consequence of environmental or spatial gradients – might affect the comparability of biomarker analyses.

PLFA analyses are often applied in soil, since they allow characterization of the complete microbiome. However, besides PLFA analysis, ergosterol is another molecule that is often used as a fungal biomarker. More data exist on quantitative ergosterol contents in fungal tissues than on PLFA biomarkers, leading to more robust conversion factors (Joergensen and Wichern, 2008 and references therein). However, the limitations discussed for PLFA data apply to an even greater extent: ergosterol is a major sterol of cell membranes in higher fungi (Moore et al., 2021), but many other sterols are incorporated in fungi and some fungi do not even produce ergosterol (Weete et al., 2010). Different fungal lineages produce specific sterol components, which again results in high variability in ergosterol concentrations among species and phyla (Joergensen and Wichern, 2008; Weete et al., 2010; conversion factors summarized by Joergensen & Wichern, 2008 range from 13 to 410 g fungal C g^−1^ ergosterol).

Due to progress in DNA-based soil analyses, quantitative PCR (qPCR) has emerged as another promising approach for the quantification of microbial groups in soil, e.g. targeting bacterial 16S rRNA genes or fungal ITS1 or ITS2 (internal transcribed spacer) sequences in the rDNA region (Fierer et al., 2005; Hungate et al., 2015; Chen et al., 2023). Currently there is only insufficient data available to conclude from qPCR results on the actual fungal biomass C. Ideally this will be possible by establishing the principles of conversion factors discussed above for PFLA and ergosterol data. Direct relationships of qPCR values and fungal biomass have yet rarely been assessed. Baldrian et al. (2013) analyzed ITS copy numbers in fungal mushrooms, reporting a conversion factor of 1.26 pg fungal C copy^−1^ ITS, with a variability slightly higher than that observed in PLFA and ergosterol contents in the same fungi. However, these analyses were done with Basidiomycete mushrooms only, where it is also unclear how active the mycelium would be compared to soil hyphae (i.e., lower DNA contents), restricted to a forest (ectomycorrhizal) system. Generally, there is an assumption that the large variability in ITS copy numbers and also mycelial DNA contents among species and environmental conditions will lead to unmanageable variation (Lofgren et al., 2019; Lavrinienko et al., 2021). However, as discussed above, other biomarkers also show large variability, with likely even stronger phylogenetic biases. In fact, qPCR values partly correlate with classical biomarker analyses (Zhang et al., 2017; Pérez-Guzmán et al., 2021; Osburn et al., 2022), and are commonly combined with them as a quantitative tool to determine group abundances (Chen et al., 2023). Thus, the growing availability of these data and their robust affiliation to respective microbial groups call for exploring the validity of qPCR conversion factors to make these data applicable for C cycling models.

Here, we aimed to provide a better understanding about fungal biomarker contents in fungi and their interspecific and phylogenetic variability to improve existing conversion factors, and explore the possibility of establishing a conversion factor for fungal ITS1 qPCR data. Based on a collection of 30 soil saprobic fungal isolates originating from natural grassland soils, covering the phyla Basidiomycota, Ascomycota, Mortierellomycota and Mucoromycota, we measured PLFA profiles, ergosterol contents and DNA concentration and ITS1 copy numbers for these isolates. These data were used to address the hypotheses that (i) high interspecific variability is present in all biomarkers, (ii) phylogenetic biases are especially strong in PLFA and ergosterol contents and (iii) high species diversity in soil can buffer against inaccuracies of conversion factors.

## Materials and methods

### Fungal isolates

A total of 30 fungal isolates of the *Rillig Lab Coreset* (RLCS) were used for this study (Andrade-Linares et al., 2016; Camenzind et al., 2022). Originally, these strains were isolated from a natural grassland site in northern Germany in 2014 (“Oderhänge Mallnow” close to the town of Lebus, Germany; 52°28′N, 14°29′E), using diverse growth media to reduce the predominance of spores and fast-growing fungi, and to include Basidiomycota strains (Andrade-Linares et al., 2016). This effort resulted in a phylogenetically diverse collection of saprobic fungi including 20 Ascomycota, 3 Basidiomycota, 2 Mucoromycota and 5 Mortierellomycota (see Table S1, Fig. 1). Still, as in all fungal studies based on direct soil isolation, the abundance of Basidiomycota was comparatively low (Thorn et al., 1996; Klamer and Bååth, 2004; Tedersoo et al., 2022).

**Fig. 1.**
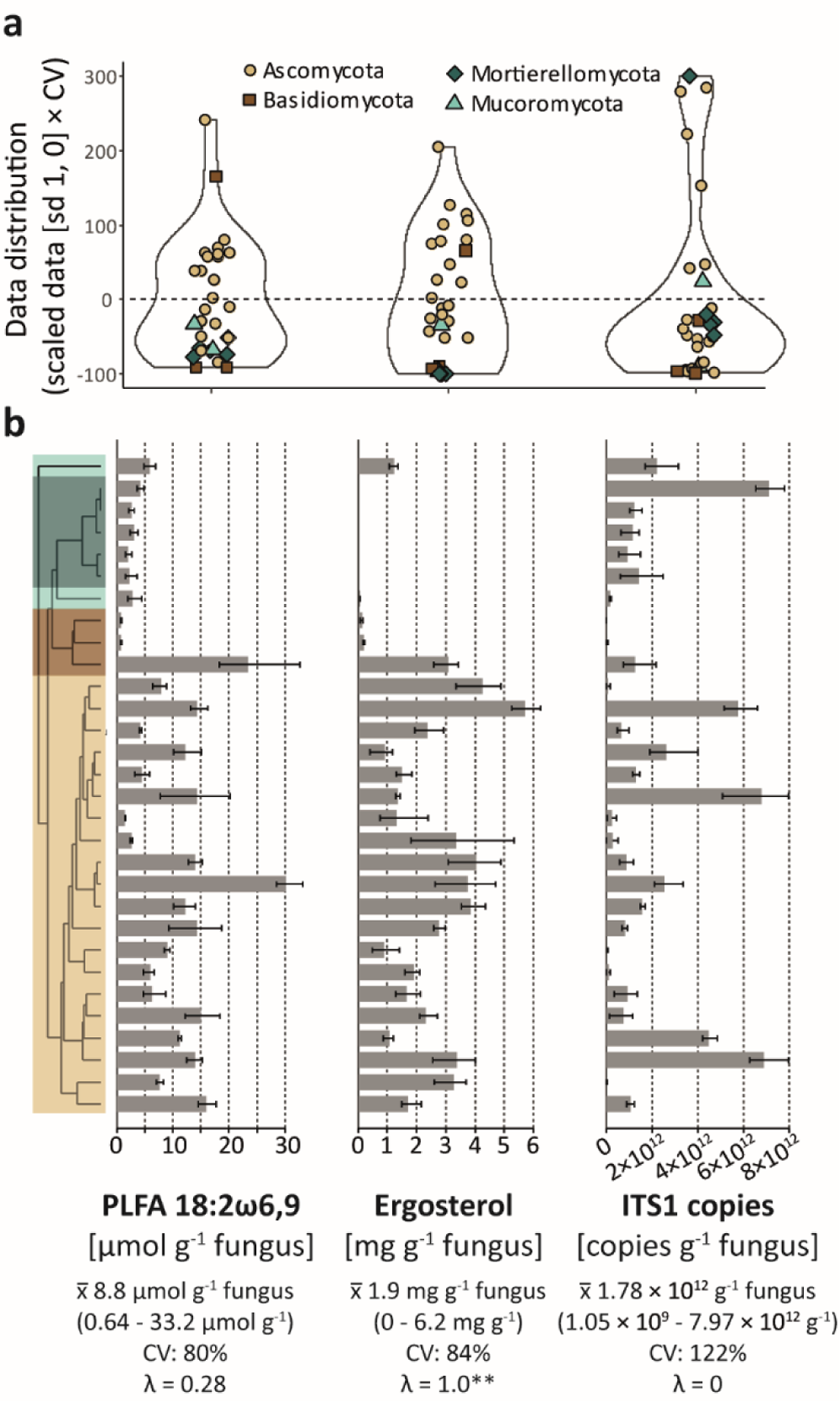
Distribution of different quantitative fungal biomarkers in soil fungal isolates. **a** Visualization of the data distribution, with the y axis representing scaled average values of each fungal isolate [1 sd, center = 0] multiplied by the coefficient of variation (CV) of each variable. This method allows for a comparison of data variability among different variables (*i.e*., a value with 1sd deviation from the mean is normalized by its proportional difference to the mean, *e.g*. 80% in case of the PLFA marker 18:2ω6,9). **b** Mean biomarker concentrations of each fungal isolate (sorted by a phylogenetic tree, colours as given in the legend above). Error bars display the minimum and maximum values (n = 3). Mean values (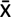), CV and Pageĺs λ of each variable are displayed below the graphs (** *P* < 0.01).

The fungal isolates have been previously characterized by functional trait analyses (e.g., Lehmann et al., 2019; Zheng et al., 2020). To maintain isolates over time, they were kept on potato-dextrose agar (PDA) at 4°C, and regularly refreshed from frozen stock cultures.

### Study design

Fungi were transferred from active PDA cultures to malt extract broth (Carl Roth GmbH + Co. KG, Karlsruhe Germany), using three repetitions per isolate. Standard growth media were needed to obtain sufficient fungal biomass for further analyses of biochemical composition, though we decided for a growth medium which is more similar to soil elemental ratios (Mouginot et al., 2014). Liquid media were chosen for simple biomass extractions without heat treatments (as commonly applied for melting agar media). To avoid artificial submersion and oxygen limitation of fungi, cultures were not shaken and kept aslant to increase accessible medium surface areas. After two weeks of growth at 20°C, fungal biomass was captured on a 20 µm mesh, and washed thoroughly (> 1 L dest. H_2_O) to remove growth media. Fungal mycelia were freeze-dried and stored at −20°C for all further analyses. Fungal growth experiments were conducted at the Freie Universität Berlin. Thereafter, samples were transferred to the University of Hohenheim for biochemical analyses. Total fungal biomass obtained ranged between 5 and 290 mg per sample, with large variations among isolates (which are known to vary strongly in growth traits (Lehmann et al., 2019)).

### Fatty acid analyses

PLFA analyses were carried out according to Frostegård et al. (1991, 1993). Initially, 2 – 20 mg fungal mycelia (depending on the available amount of fungi) were lysed with 1 ml Bligh & Dyer reagent in a Lysing Matrix E Tube (MPBio) using a FastPrep Instrument (5,5 Speed, 40 s) and then mixed with 17.4 ml Bligh & Dyer reagent [chloroform, methanol, and citrate buffer (pH 4), 1:2:0.8]. For lipid fractionation, the extract was pipetted onto a silica acid column and subsequently eluted from the column with chloroform (NLFA), acetone (Glycolipids), and methanol (PLFA) (Frostegård et al., 1991). Methanolysis of the PLFA fraction was conducted with methanolic KOH and fatty acid methyl esters (FAMEs) were extracted with a solution of hexane chloroform and acetic acid as described by Ruess et al. (2007). The resulting PLFA were dissolved in iso-octane and measured on a gas chromatograph (GC, AutoSystem XL, PerkinElmer Inc., Norwalk, CT, United States). The GC was equipped with a flame ionization detector, an HP-5 capillary column (crosslinked 5% phenyl methyl silicone; 50 m x 0.2 mm, film thickness: 0.33 mm) and helium as carrier gas. The initial column temperature of 70°C was held for 2 min. Temperature was then increased by 30°C min^−1^ to 160°C, then by 3°C min-1 to 280°C and held for 15 min. Injection temperature was set at 260°C. The concentration of FAMEs was calculated via an internal c24:1 FAME standard, which was added to the samples before methylation.

### Ergosterol contents

Ergosterol was determined according to Djajakirana et al. (1996). In brief, 1 - 13 mg fungal mycelia (depending on the available amount of fungi) was weighed into a Lysing Matrix E Tube (MPBio), 1 ml of ethanol was added and lysed in the FastPrep like for PLFA analysis. The lysate was transferred into a 100 ml amber glass wide-neck bottles and extracted with all in all 25 ml ethanol on a horizontal shaker for 30 min at 250 rpm. After extraction, the suspensions were transferred to 50 ml Falcon tubes and centrifuged for 30 min at 4400 *x g*. Ten ml of the extracts were then transferred to 10 ml centrifuge tubes and vaporized to dryness in a rotary vacuum concentrator at 50°C. Subsequently, the dried extracts were dissolved in 1 ml methanol and transferred through syringe filters (0.45 μm) into amber glass HPLC vials. For each extraction, two standard soil samples as well as blanks (without soil) were included. The measurements were performed via HPLC (Agilent 1260 Infinity series, Agilent Technologies). For ergosterol separation, a reversed phase column (MZ Spherisorb ODS-2 C18, 150 x 3 mm, particle size 3 µm) and 100 % methanol as mobile phase (flow rate of 0.5 ml min^−1^) was used. Temperature during the measurement was set to 40°C. The detection occurs with a Diode Array Detector (Agilent) at a wavelength of 282 nm.

### DNA concentrations and ITS1 copy number determination

The abundance of the fungal ITS1 DNA sequence was quantified by qPCR (ABI prism 7500 Fast System, Applied Biosystems, Foster City, CA, USA). For this, DNA was extracted from 1 - 13 mg fungal mycelia (depending on the available amount) using the E.Z.N.A. Soil DNA Kit (Omega Bio-Tek, Inc., VWR International GmbH, Bruchsal) and quantified with a Picogreen kit (Quant-IT^TM^ PicoGreen^TM^ dsDNA Assay kit, Invitrogen) according to the protocol of the company. DNA was diluted with ultra-pure water to a target concentration of 3 ng DNA μl^−1^. For the qPCR assay a reaction mix of 0.75 μl of each forward (ITS3F) and reverse (ITS4R) primer (White et al., 1990; Manerkar et al., 2008; primer concentrations of 10 pmol μl-1), 0.375 μL T4gp32, 7.5 μl SYBR Green, 4.125 μl ultra-pure water, and 1.5 μl DNA template was prepared. A program with 35 cycles, each 10 min 95°C, 15 s 95°C, 30 s 55°C, 30s 72°C, 30s 76°C was used. Additionally, a 60°C to 95°C step was added to each run to obtain the denaturation curve specific for each amplified sequence. Mean extraction efficiency was 93%. Standard curves were generated in triplicate with serial dilutions of a known quantity of the respective isolated plasmid DNA. Each qPCR run included two no-template controls showing no or negligible values.

### Calculation of conversion factors

Following the methods established by Joergensen and Wichern (2008), we calculated conversion factors for the commonly used quantitative soil fungal biomass estimates - 18:2ω6,9 PLFA marker [mg fungal C µmol^−1^ 18:2ω6,9], ergosterol [g fungal C g^−1^ ergosterol] and ITS1 copy numbers [pg fungal C copy^−1^ ITS]. We calculated conversion factors based on the dataset presented here, as well as on available data from a literature review, describing biomarker contents in fungal tissues (for the complete dataset see Table S2). In case studies assessed biomarker contents for different replicates, time points or conditions, either mean values were taken or only one treatment included, to obtain one value per fungal strain (complete datasets are summarized in Table S2). For the calculation of conversion factors, mean biomarker contents in all tested fungal isolates (in this study or all studies, respectively) were taken, and corrected for the relative C content in fungal biomass – here estimated as 0.47. This value was also used by Joergensen and Wichern (2008), and perfectly coincides with the average C content in the fungal isolates analyzed here when grown under different growth media conditions (10% potato-dextrose agar, cellulose medium and glucose medium with C:N 20; Camenzind et al., 2021). In case of the PLFA marker 18:2ω6,9 an additional phylogenetically corrected conversion factor was calculated, due to the large variation in marker contents among phyla. In this case, a weighted mean of PLFA marker contents in fungal isolates was calculated using global read abundances of the four phyla reported in Tedersoo et al. (2022; Table S2).

### Statistical analyses

All statistical analyses were conducted in R version 4.1.3 (R Core Team, 2021). Respective commands and packages are given in brackets.

The phylogenetic relatedness among fungal isolates was determined using an alignment of the full sequence reads of ITS1, 5.8S, ITS2 and partial LSU (AlignSeqs(), *DECIPHER* (Wright, 2016)). Genetic distance was calculated (dist.ml(), *phangorn* (Schliep, 2010)) and a phylogenetic tree constructed using the unweighted pair group method with arithmetic mean (upgma(), *phangorn*).

The mean (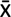), coefficient of variation (CV = (standard deviation / mean) * 100) and phylogenetic signal (Pageĺs λ; phylosignal(), package *phylosignal* (Keck et al., 2016)) were calculated for each variable based on the average values of individual isolates (*n* = 3). Differences among phyla were analyzed with the whole dataset (all replicates included), using mixed-effects models with isolate identity as random effect (lme(), *nlme* (Pinheiro et al., 2021)). In case of non-random distributions of residuals, generalized linear mixed models were fit with appropriate error distributions (glmmPQL(), *MASS* (Venables and Ripley, 2002)). PLFA patterns were displayed by principal component analyses (PCA, prcomp()) using relative PLFA marker contents.

We modeled the effects of interspecific variability in marker contents on the validity of conversion factors based on theoretical fungal soil communities. To analyze the overall effects of community shifts on deviations of estimated fungal C contents (calculated using isolate biomarker contents and our conversion factors) from actual fungal C contents, randomized communities of different diversity levels of the 30 fungal isolates analyzed here were modeled (with the assumption that biomarker expression does not change in a community context). For each diversity level (5, 10, 20 and 30), 1000 random communities were generated based on random draws (without replacement) of fungal isolates. Thereafter, individual isolate abundances in these theoretical communities were randomly varied between 0 and 100. For each of these randomized communities, the real fungal C content (fungal C = 47% of biomass) based on summed fungal abundances was calculated. The estimated fungal C content was calculated summing biomarker contents of included isolates, which were converted to fungal C with conversion factors obtained in this study (here we used the conversion factors calculated with our data only).

Additionally, the validity of the conversion factors was modeled in soils differing in the composition of fungal phyla. We chose to base these model communities on three exemplary soils used in a common experimental platform of the SoilSystems project (DFG project SPP 2322, www.soilsystems.net), which strongly varied in their phylogenetic composition. Details on fungal community analyses in these soils are summarized in the Supporting Information S1. Shortly, soil fungal abundance was quantified by qPCR, and community composition determined by Illumina sequencing of the ITS1 taxonomic marker. These data revealed a strong variation in the relative abundances of fungal phyla. Here, we paired the information on total fungal abundance and relative phyla variation to model theoretical communities with our 30 fungal isolates. Again, for each modeled soil type, 1000 random communities were assembled (each based on all 30 fungal isolates). Within these communities, abundances of individual isolates were varied randomly, though with the boundaries of relative phyla abundances as observed in these soil types. Total fungal C contents were based on the relative fungal ITS copy numbers observed in these soils (Supporting Information S1). As described above, biomarker based fungal C contents were estimated using PLFA biomarker contents in modeled communities and respective conversion factors, and compared to actual fungal C contents.

## Results

### Expression of soil fungal biomarkers in saprobic fungal isolates

On average, the fungal isolates tested in this study contained per g fungus (dry weight) 1.87 ± 1.6 mg ergosterol (mean ± s.d.), 1.78 × 10^12^ ± 2.2 × 10^12^ ITS1 copy numbers and 173 ± 113 µg total DNA contents (Fig. 1). The total PLFA concentration was on average 22.0 ± 11 µmol g^−1^ fungus, with the highest peak being present in the fungal biomarker 18:2ω6,9 with 8.8 ± 7 µmol g^−1^ fungus, as well as the 18:1ω9c/18:3ω3,9,12 and 18:3ω6,9,12 markers with 4.8 ± 3 and 0.55 ± 1.6 µmol g^−1^ fungus, respectively (Table 1, Fig. 2).

**Fig. 2.**
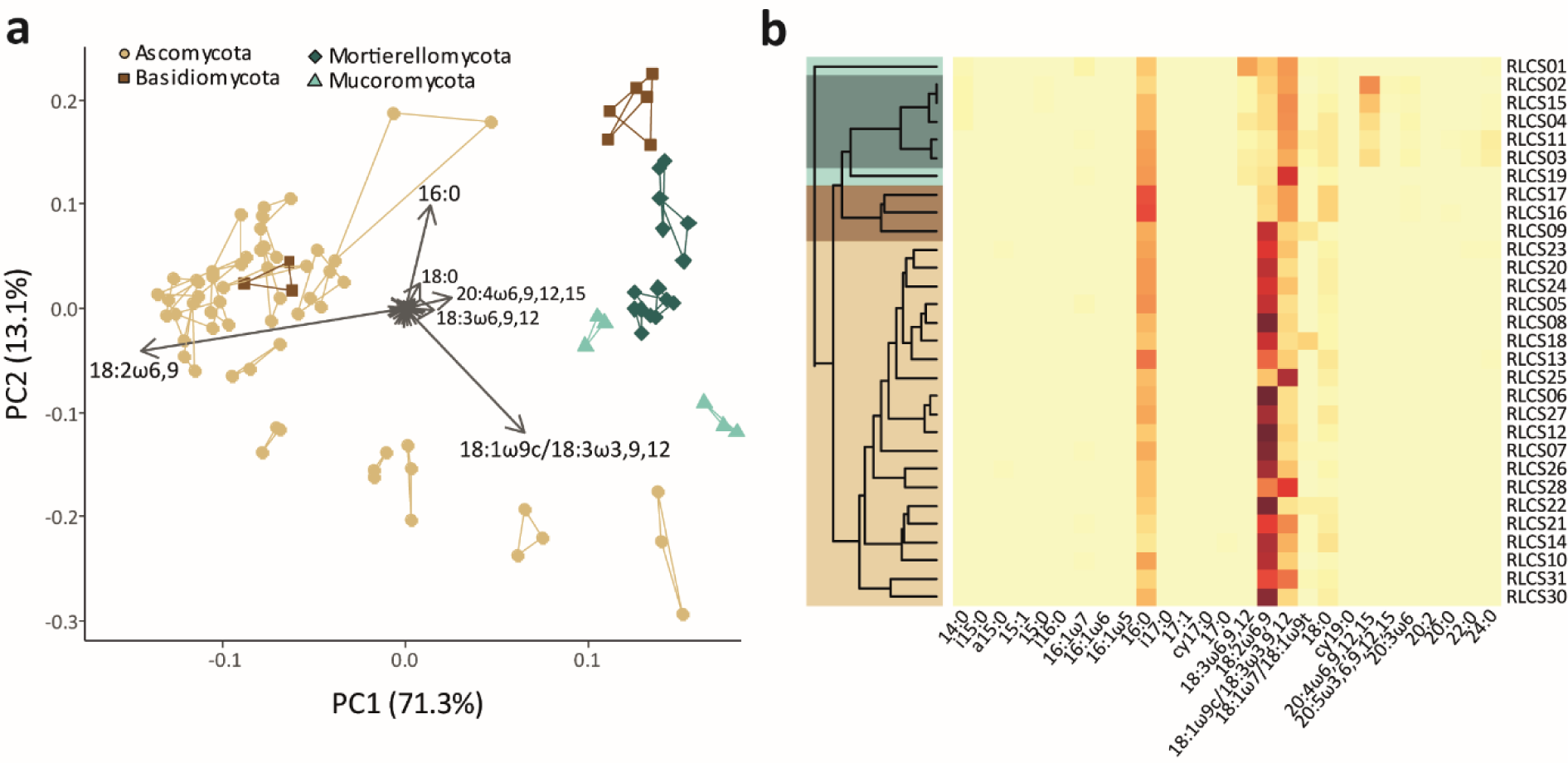
Visualization of the PLFA marker distribution in different soil fungal isolates. **a** Principal component analysis (PCA) of the relative abundance of different PLFAs. Dot shape and color reflect respective phyla, dots connected by lines are repetitions of the same isolate. **b** Heatmap of the relative abundances of individual PLFAs in fungal isolates.

**Table 1.**
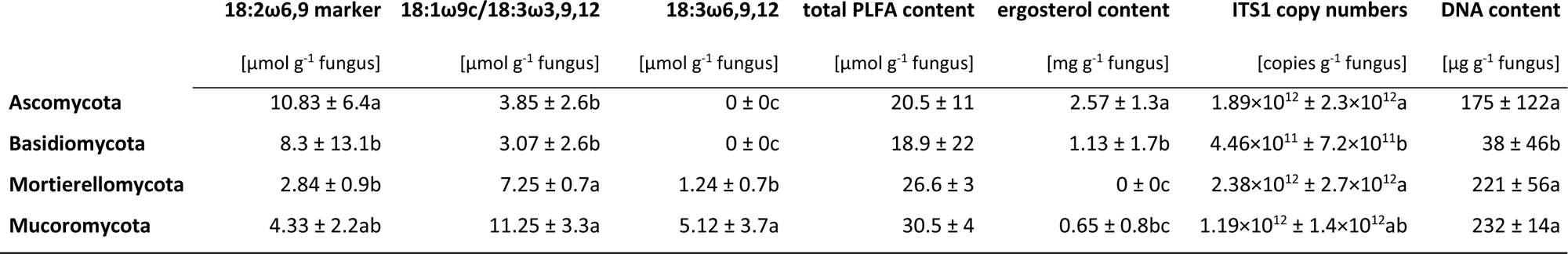
Fungal biomarker distribution in different fungal phyla, with mean values ± standard deviations displayed (letters indicate significant differences among phyla; values are based on 30 isolates, 3 repetitions each).

There was considerable variation in biomarker contents among fungal isolates, as also evident from the large standard deviations and coefficients of variation (Fig. 1b). The overall interspecific variation was highest in ITS1 copy numbers (CV = 122 %, Fig. 1). However, this ITS1 copy number variation was not driven by phylogenetic relationships, whereas ergosterol contents and fungal PLFA markers showed a clear deviation among phyla (Table 1, Fig. 1), reflected also in the phylogenetic signals (Fig. 1b). When analyzing differences among the individual phyla, there was a clear difference in Mortierellomycota and Mucoromycota compared to Ascomycota and Basidiomycota (Fig. 2a). Both the concentrations of the fungal PLFA biomarker 18:2ω6,9 and ergosterol were significantly reduced (Fig. S1a), while the PLFA markers 18:1ω9c/18:3ω3,9,12 and 18:3ω6,9,12 were enriched. The PLFA marker 18:3ω6,9,12 was even only present in these two groups (Table 1), while PLFA 20:4ω6,9,12,15 was only present in Mortierellomycota (Fig. 2b). By contrast, total PLFA contents, ITS1 copy numbers and DNA contents did not show these phylogenetic patterns. For the Basidiomycota only three isolates were available in total. Two of them (RLCS09 and RLCS17), both Agaricales, showed very low concentrations of all measured biomolecules, i.e., DNA, ergosterol and PLFAs. Since both isolates grew very well on the selected media producing sufficient biomass (data not shown), it is not clear whether this result is a methodological issue of the extraction methods, or relates to special physiology of these isolates. Results of these Basidiomycete isolates should be interpreted with caution and need further investigation for verification.

### Calculation of conversion factors for soil fungal biomarkers

Conversion factors for different biomarkers are presented in Table 2, based on biomarker contents in fungal isolates analyzed in this study (see above), as well as on mean values including further available data (Table S2).

**Table 2.** Conversion factors based on our dataset, and a literature review of available data on fungal isolates (data given in Table S2).

| Soil fungal biomarker | Remarks | Unit | Conversion factor<br>(our data) | Average conversion factor<sup>2</sup><br>(including further datasets) | # studies<br>included |
| --- | --- | --- | --- | --- | --- |
| Ergosterol<sup>1</sup> | Reflects the abundance of Ascomycota and Basidiomycota only | [g fungal C g<sup>-1</sup> ergosterol] | 197.5 (CV 60%<sup>1</sup>) | 95.34 (mean CV 46%) | 13 |
| PLFA marker 18:2ω6,9 | Phylogenetic correction needed in case phyla vary among soils | [mg fungal C μmol<sup>-1</sup> 18:2ω6,9) | 53.3 (CV 80%) | 32.0 (mean CV 65%) | 3 |
| PLFA marker 18:2ω6,9 - phylogenetically corrected | Correction based on read abundances of phyla given in Tedersoo et al. 2022 | [mg fungal C μmol<sup>-1</sup> PLFA 18:2ω6,9) | 53.0 | 26.8 | 3 |
| ITS1 copy numbers | Method-specific variations | [pg fungal C copy<sup>-1</sup> ITS1] | 0.264 (CV 122%) | n.a. |  |

<sup>1</sup>data from Mucoromycota and Mortierellomycota were excluded in these calculations

The conversion factor for the PLFA biomarker 18:2ω6,9 amounted to 53.3 mg fungal C µmol^−1^ 18:2ω6,9 based on data presented in this study. Contents of the PLFA biomarker 18:2ω6,9 in fungal isolates were further available from two additional studies (to the best of our knowledge). Klamer and Bååth (2004) used very similar methods as applied here with a comparable phylogenetic coverage, revealing similar biomarker contents in fungal tissues (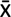 = 6.61 µmol g^−1^ fungus; Table S2). By contrast, Baldrian et al. (2013) analyzed PLFA contents in basidiomycete mushrooms, reporting average 18:2ω6,9 contents 4.5 times higher than observed in our study (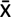 = 39.6 µmol g^−1^ fungus; Table S2). These high values were also driven by high relative contents of 18:2ω6,9 in extracted PLFAs – on average 86%, compared to 40% in our study, or 45 and 43% reported by Klamer and Bååth (2004) and Federle et al. (1986). Therefore, the average conversion factor calculated with all three datasets was significantly lower (Table 2). The estimate for a phylogenetically corrected conversion factor based on all three datasets (corrected by global read abundances) was similarly lower, driven by higher values for Basidiomycota in Baldrian et al. (2013; Table 1 and S2). The data given in Table S2 also allow to calculate corrected conversion factors for individual soils.

In case of ergosterol we included 12 additional studies assessing ergosterol contents in fungal tissues. Since ergosterol was not (or only in low amounts) observed in Mortierellomycota and Mucoromycota (Fig. 1), we excluded these phyla from the calculation of all conversion factors for ergosterol (Table 2). The average fungal ergosterol content reported in individual studies varied between 0.21 and 15.23 mg g^−1^ fungus, with an overall mean value of 4.93 mg g^−1^ fungus (Table S2). Since this relates to more than double the average ergosterol content in the isolates analyzed here (Asco- and Basidiomycota only), the average conversion factor including all studies was significantly lower (Table 2).

Based on the average ITS1 copy numbers in fungal isolates analyzed in this study (1.78 x 10^12^ g^−1^ fungus, Fig. 1), we also calculated a first proxy for a conversion factor for ITS1 copy numbers: 0.264 pg fungal C copy^−1^ ITS1 (Table 2). Few other studies calculated ITS1 copy numbers in fungal tissues, but the applied methods varied strongly making direct comparisons challenging (Table S2). Therefore, no average conversion factor was calculated in this case. Other studies not only applied different growth conditions (affecting the age of the mycelium and DNA contents), but also different primer pairs, DNA isolation techniques and qPCR methods (Table S2). The resulting mean values of ITS1 copy numbers in fungal tissues showed a wide range: 1.88 x 10^09^ - 1.27 x 10^12^ g^−1^ fungus. Only one study also included several isolates to quantify ITS copy numbers (Baldrian *et al*., 2013), using mushroom tissues of Basidiomycota only. These analyses resulted in slightly lower ITS copy numbers than presented here (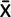 = 3.74 x 10^11^ g^−1^ fungus), and less variation among fungal isolates (CV = 78%).

### Modeling the effects of interspecific variability on the validity of conversion factors

As shown above, biomarker contents varied strongly among fungal isolates. The conversion factors represent an average value based on a range of different isolates, and thus may correctly reflect the overall C content in a community with equal abundances of these isolates. However, in case of abundance shifts among community members, fungal marker contents (relative to fungal C contents) will shift in parallel, resulting in an under- or overestimation of fungal C contents based on biomarker analyses (Fig. 3). In modeled fungal communities with randomly varied abundances, the PLFA biomarker overall revealed more robust estimates of fungal C contents than ITS copy numbers (Fig. 3). At a diversity level of 30 isolates, 95% of the calculated fungal C estimates deviated between −16% to +16% from actual fungal C contents in case of the PLFA biomarker, while with ITS copy numbers deviations ranged between −26% and 24%. The diversity level had a strong effect on these numbers. If only few isolates were present in a modeled community, shifts in their abundances affected the validity of biomarker estimates much more strongly than in communities with more fungal isolates. At the lowest diversity level of only 5 isolates, the deviation of fungal C estimates from actual C contents varied between −68% to 87% (95% data range) for the PLFA biomarker, and between −78% and 127% for the ITS copy numbers (Fig. 3).

**Fig. 3.**
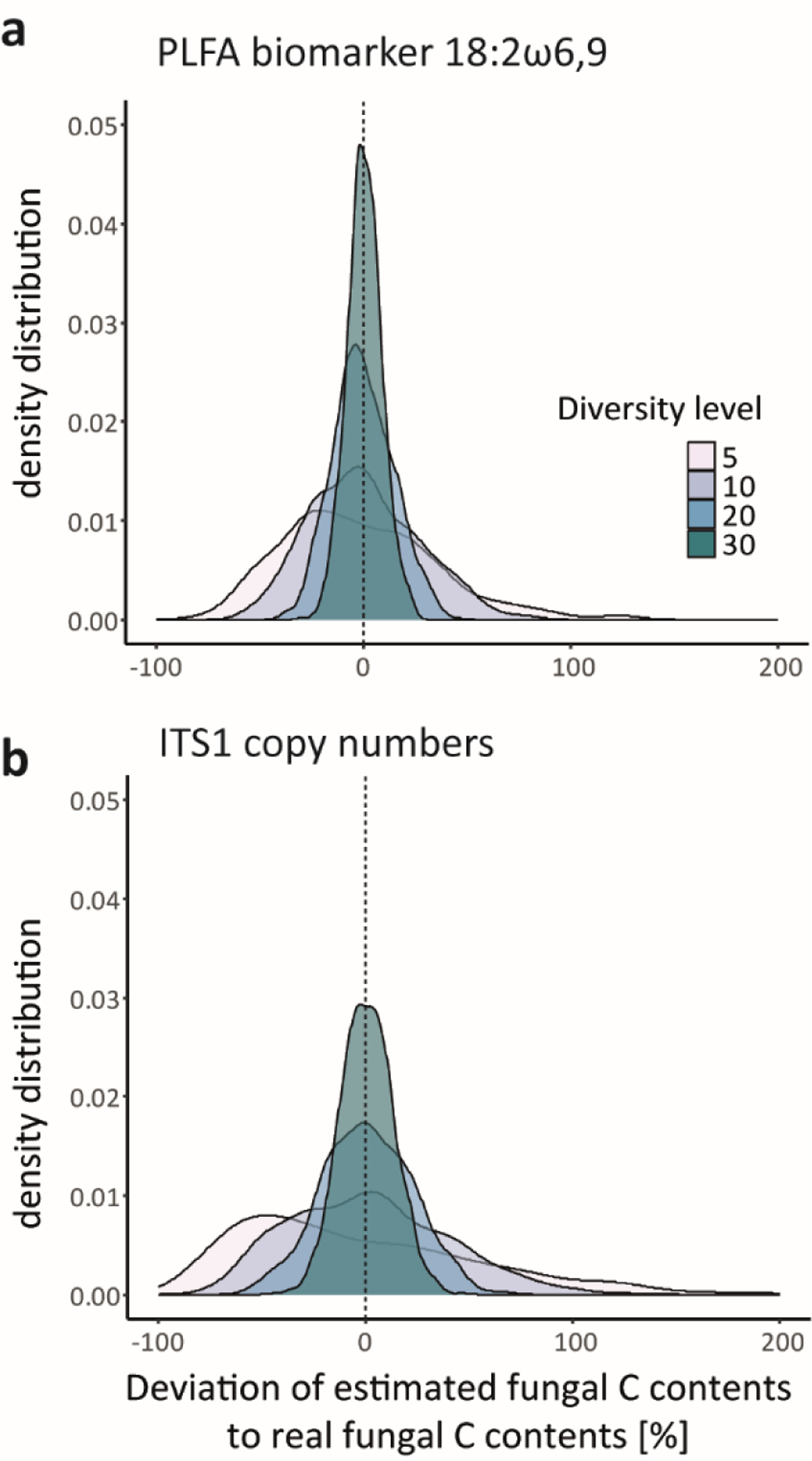
Modeling outcome of the effects of fungal community shifts on the validity of conversion factors to estimate soil fungal C contents under different diversity levels. Density distributions of the deviation of biomarker fungal C estimates from actual fungal C contents based on modeled communities are displayed, based on the PLFA biomarker 18:2w6,9 (**a**) and ITS1 copy numbers (**b**). Randomized abundances of fungal communities with 30 isolates were modeled at different diversity levels (1000 runs for each diversity level). For each community, the fungal C content was calculated based on biomarker contents with conversion factors. These values were compared with the actual fungal C contents present in each community. Individual colors visualize different diversity levels.

In theoretical communities of soils with different phylogenetic composition, the estimated fungal C contents in modeled fungal communities partly diverged from the actual relative fungal C contents (Fig. 4). All three soil types had relatively high abundances of Mortierellomycota (Fig. 4; Tedersoo et al., 2022), which led to an underestimation of fungal C contents when applying our conversion factor for the PLFA biomarker (Fig. 4a). This underestimation was stronger in soil types with lower relative abundances of Ascomycota. The relative difference in fungal C contents among soils was captured correctly when considering only the interquartile range of the modeled communities. In certain cases of random community composition, though, the difference in fungal abundances among soils 1 and 3 would have not been detected based on PLFA biomarker data (Fig 4a). In the case of the ITS1 marker, high relative abundances of Mortierellomycota and low abundances of Basidiomycota led to a slight overestimation of actual fungal C contents in all three soils (Fig. 4b). This deviation was less pronounced than in the PLFA biomarker, however, overall variability in fungal C estimates was higher.

**Fig. 4.**
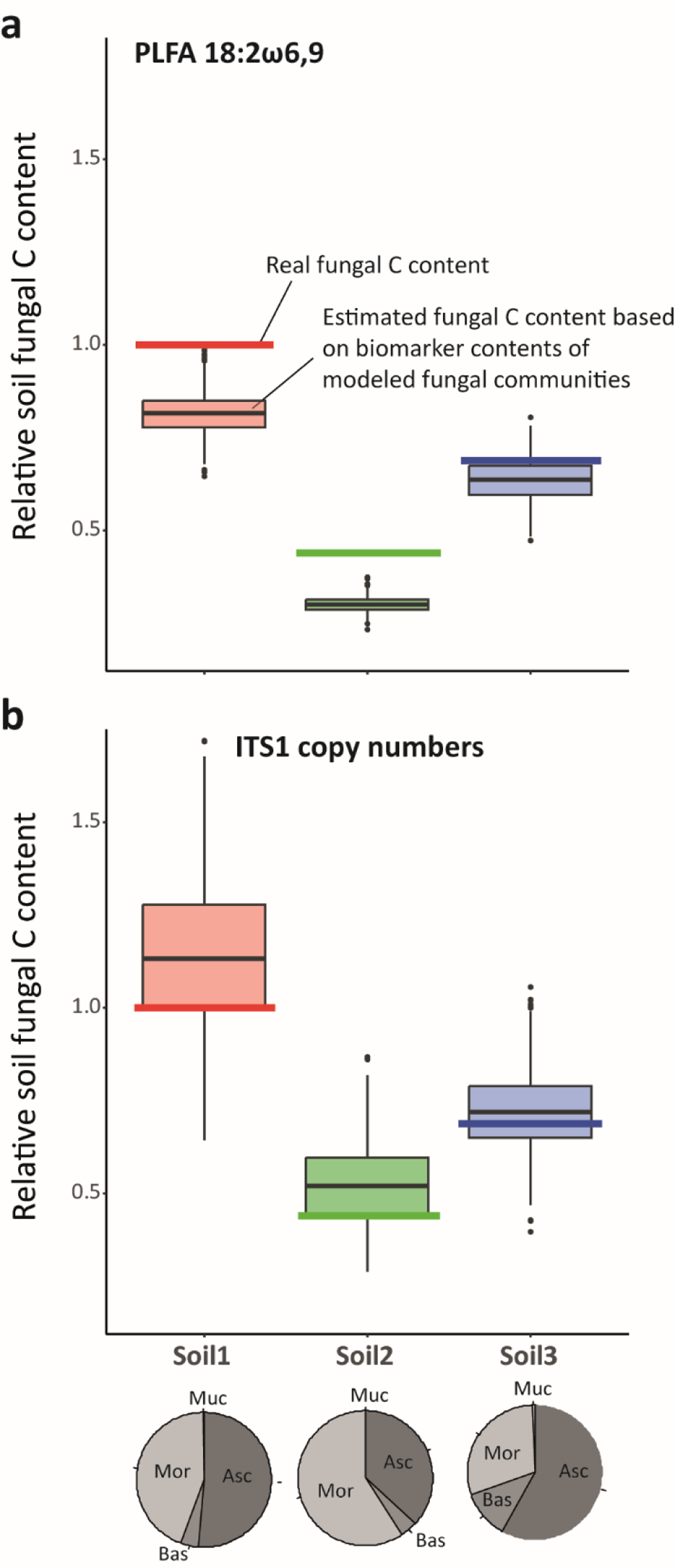
Modeling outcome with theoretical composition of fungal isolates according to real phyla distribution and fungal quantity in different soils (for details on selected soils see Supporting Information S1), applying conversion factors for (**a**) PLFA 18:2w6,9 and (**b**) ITS1 copy numbers. Solid lines represent the actual relative fungal C contents in three different soils, boxplots depict the estimated fungal C contents based on biomarker contents in modeled communities of 30 fungal isolates (colors distinguish the three soil types). Fungal abundances were randomly varied within the boundaries of the relative phyla distribution in these soils. Pie-charts were added to depict these relative abundances based on Illumina sequencing reads (Supporting Information S1).

## Discussion

Biomarker analyses in a collection of fungal isolates provided novel conversion factors and revealed the extent and consequences of interspecific variability for their application in soil. Observed fungal PLFA biomarker and ergosterol contents were within the range of other laboratory fungal analyses, adding to the refinement of existing biomarker conversion factors (Klamer and Bååth, 2004). Additionally, data on fungal ITS1 copy contents provided a first estimate of a conversion factor for fungal qPCR data, though interspecific patterns and comparisons with other data also highlight pros and cons for qPCR application in soil. As hypothesized, interspecific variability was high in all biomarkers. Though in fact, it was most pronounced in ITS1 copies g^−1^ fungus, also when analyzing other (published) datasets. On the other hand, in support of our second hypothesis, we detected a stronger phylogenetic bias in the fungal PLFA biomarker and ergosterol than in ITS1 copies, mostly driven by differential marker profiles in Mucoromycota and Mortierellomycota. A modeling exercise gave first insights into the buffering capacity of fungal diversity in soil for conversion factor accuracy: as hypothesized, higher levels of fungal diversity in soil have the potential to reduce biases introduced by high species variability, whereas phylogenetic shifts in fungal communities might affect the validity of biomarker comparisons among soils, especially for the fungal PLFA biomarker (and ergosterol; not explicitly tested here).

### Fungal PLFA profiles and ergosterol contents as biomarkers

Analyses of fungal PLFA profiles supported the existence of phylum specific markers, although the lack of specificity hampers its full application potential in soil. We focused in this study on the fungal PLFA biomarker 18:2ω6,9, which is widely applied to estimate fungal abundances and fungal:bacterial ratios in soil. As suggested in previous studies (Federle, 1986; Zelles, 1997; Baldrian et al., 2013), 18:2ω6,9 was overall the most abundant PLFA marker in fungal isolates tested here. This marker is commonly used as a fungal biomarker due its high concentrations in fungal biomass (as confirmed here), but also due to its specificity for fungi in soil microbial communities (Ruess and Chamberlain, 2010; Joergensen, 2022). The only bias may be given by plant tissue, i.e., roots, which Joergensen (2022) suggests to reduce by picking them before soil PLFA extraction. Other PLFA markers enriched in fungi are not uniquely found across groups (Ruess and Chamberlain, 2010), though they would add improved phylogenetic coverage of all fungal lineages. 18:2ω6,9 is only dominant in Ascomycota and Basidiomycota, while Mortierellomycota and Mucoromycota have different PLFA profiles: the presence of 18:3ω6,9,12 in fungal isolates (even though in low concentrations) in these phyla confirmed this marker as a “Zygomycete peak” (Weete, 1980; Van der Westhuizen et al., 1994), similar to the patterns of 18:1ω9c (also suggested to be found in “Zygomycetes”) and 18:3ω3,9,12 (Basidiomycota and Ascomycota; Ruess and Chamberlain, 2010; Joergensen, 2022; though unfortunately the latter two were only co-eluted in this study). Additionally, we detected a high concentration of the PLFA 20:4ω6,9,12,15 in *Mortierella* isolates, which was also reported by Nisha et al. (2011). However, a- and g-linolenic acid as well as arachidonic acid are also abundant in other microbial groups, *i.e*., bacteria and algae, microfauna and plant debris (Ruess and Chamberlain, 2010; Frostegard et al., 2011; Willers et al., 2015). For these reasons, the fungal biomarker 18:2ω6,9 is most specific to describe fungal abundance in soil, and thus demands a conversion factor (Joergensen and Wichern, 2008).

The PLFA analyses presented here add important data points to inform this PLFA 18:2w6,9 conversion factor. So far, insufficient data existed to translate PLFA biomarker values to fungal C contents in soil. The conversion factor based on fungal isolates analyzed here (53.3 mg fungal C µmol^−1^ 18:2ω6,9 PLFA) was within the range of previously reported conversion factors (Klamer and Bååth, 2004: 85 mg fungal C µmol^−1^ 18:2ω6,9 PLFA; Joergensen and Wichern, 2008 based on additional correlations with soil hyphal length: 107 mg fungal C µmol^−1^ 18:2ω6,9 PLFA; Baldrian et al., 2013: 12 mg fungal C µmol^−1^ 18:2ω6,9 PLFA). Since conversion factors vary notably between studies (as well as within selected isolates) we propose an average conversion factor based on all available data: 32.0 mg fungal C µmol^−1^ 18:2ω6,9 PLFA. An average value reduces biases by phylogenetic coverage and selection of isolates in individual studies, and improves the general validity of the conversion factor.

The number of available reference data to calculate conversion factors varied widely among biomarkers. In case of ergosterol contents, comprehensive data are available (Joergensen and Wichern, 2008). The ergosterol contents observed in fungal isolates tested here were comparatively low, but within the range of previous reports (Table S2). Overall, 13 datasets were included for ergosterol contents, which revealed an average conversion factor of 95.34 g fungal C g^−1^ ergosterol. By comparison, only Baldrian et al. (2013) tested ITS copy contents in a meaningful number of fungal isolates (to the best of our knowledge). This lack of data impairs our understanding of ITS copies as a fungal biomarker.

### Applicability of a conversion factor for ITS1 qPCR data

Knowledge about ITS copies in fungal biomass is important for the interpretation of qPCR based fungal quantification in soil. The large deviation among fungal ITS copy contents in different studies was likely driven by methodological differences: especially differences in DNA extraction methods and primers applied (Table S2) can result in varying DNA extraction efficiencies (Delmont et al., 2012) and qPCR results, even when different primer pairs target the same gene (Thijs et al., 2017). To allow for comparability among datasets, methods of qPCR quantification of soil fungi need to be standardized, as is the case for PLFA or ergosterol analyses (Frostegård et al., 1991; Sae-Tun et al., 2020). In our study, fungal ITS1 copy contents led to a conversion factor of 0.264 pg fungal C copy^−1^ ITS1. However, the CV among isolates was high with 122% and higher than in PLFA biomarker and ergosterol contents. The large interspecific variability in ITS1 copies is likely driven by two components: copy number variation within the genome, as well as variability in total DNA concentrations (both correlated, R² = 0.17, Fig. S1b). In fungi, high rDNA copy number variation (including the ITS region) was reported based on sequenced genome analyses: in 91 fungal taxa Lofgren et al. (2019) detected a wide range of numbers: 14 to 1442 rDNA copies in the genome. In their analyses, Ascomycota had on average lower copy numbers compared to Basidiomycota and early diverging fungal lineages. On top of this variation, DNA contents also vary among isolates: in our study DNA contents showed a CV of 66% among fungal isolates (data not presented). Baldrian et al. (2013) report similar variation - CV of 62% in fungal DNA contents. Thus, the resulting high variation among isolates in derived ITS copies g^−1^ fungus is an interplay of copy number variation in genomes and variable total DNA contents.

Such high variation in ITS copies and DNA contents is not only problematic for finding valid conversion factors and for comparing fungal abundances among soils or treatments, but it is an important issue to consider in sequencing studies (Větrovský and Baldrian, 2013; Lavrinienko et al., 2021). As shown by data presented here, not only preferential primer binding to certain sequences, but also interspecific variability in ITS copy contents may bias the assignment of sequencing read numbers to actual fungal species abundances in soil.

### Accuracy of conversion factors in highly diverse soil samples

Variability in biomarker contents among fungal species is evident; but, on the other hand, microbial diversity in soil is exceptionally high (Anthony et al., 2023). This has led to the hypothesis that high species diversity in soil buffers against variability in marker contents. Interspecific variability was not only high in ITS copies, but also in PLFA biomarker and ergosterol contents in our fungal isolates (CV of 80% and 84%, respectively), similar to values detected in other datasets (Table S2). Consequently, differences in fungal species composition might impact the comparability of conversion factors applied to different soil samples (Nurika et al., 2018): a biomarker conversion factor only provides one value for the complete fungal community, even though biomarker : fungal C ratios differ among species. Variability leading to inaccurate conversion factors not only affects the correct translation of biomarker contents to fungal C, but also the validity of comparisons among soils and treatments based on biomarker analyses. Therefore, the following discussions not only concern conversion factors, but the general interpretation of soil fungal biomarker analyses.

Our modeling approach confirmed that differences in species composition affect the validity of conversion factors, even stronger for ITS1 copies due to high variability among species. However, in line with our third hypothesis, the more species present in a community, the lower the impact of interspecific variability, *i.e*., the higher the accuracy of conversion factors (and correct assignment of biomarker contents to actual fungal biomass). Indeed, this is a fundamental, universal principle often referred to as the Regression Toward the Mean (Stigler, 1997). In the present case, sampling a greater number of diverse taxa with inherently different ergosterol, PLFA or ITS1 copies will cause these measurable values to converge upon the true mean. Even though these conclusions are based on the modeling of 30 isolates only, we think these principles also apply *in situ* to soil fungal communities. The range of biomarker values in our fungal isolates was comparable to other studies (Barajas-Aceves et al., 2002; Klamer and Bååth, 2004; Baldrian et al., 2013), and broadly covered diverse fungal lineages. In case fungal marker contents are comparable in soil (a prerequisite for the application of these biomarkers in general, see discussion below), the even larger diversity of fungal communities in soil samples should only reinforce the accuracy of conversion factors and comparability of biomarker analyses (Tedersoo et al., 2020).

Beside interspecific variability, also phylogenetic biases are problematic in soil biomarker application. Differences among phyla were present in fungal PLFA biomarker and ergosterol contents. Ergosterol generally only represents Ascomycota and Basidiomycota (Weete et al., 2010), as confirmed by our data. Similarly, the PLFA biomarker was significantly reduced in Mortierellomycota and Mucoromycota. This difference among phyla in PLFA biomarker contents indeed led to an underestimation of fungal C contents in soils characterized by high relative abundances of these phyla, when applying the proposed conversion factor in theoretical models. Especially in incubation experiments with glucose this bias must be considered, since Mucoromycota and Mortierelloymycota represent early-successional “sugar fungi” (Kramer et al., 2016; Pawłowska et al., 2019). By contrast, there was no phylogenetic signal detected in ITS1 copies g^−1^ fungus, thus, accuracy of this conversion factor was comparable among soils despite contrasting abundances of fungal phyla. The agricultural soils used here as a modeling template were characterized by high read abundances of Mortierellomycota. Tedersoo et al. (2022) report much lower relative sequence abundances for this phylum in a global comparison (an average 8% of reads for Mortierellomycota, 3% Mucoromycota), though agricultural management and environmental factors can indeed increase their relative abundances (Wang et al., 2020; Li et al., 2023). These groups are often overlooked as early diverging fungal lineages, but they include relevant and abundant saprobic species in soil and should be covered by fungal C estimates (Pawłowska et al., 2019; Tedersoo et al., 2022). Thus, regarding phylogenetic coverage and comparability, ITS copies represent a more suitable fungal biomarker. Differences among Ascomycota and Basidiomycota cannot be resolved based on fungal isolates included here (two of three Basidiomycota isolates had low extraction efficiencies for all biomarkers). In PLFA biomarker and ergosterol contents there seems to be no substantial difference among these two phyla (Antibus and Sinsabaugh, 1993; Barajas-Aceves et al., 2002; Klamer and Bååth, 2004; Baldrian et al., 2013). Potentially, reports about low rDNA copy numbers in Ascomycota suggest phylogenetic biases also in qPCR data (Lofgren et al., 2019). In an ecological context, variation in organismal rDNA is often correlated with high growth rates (Sterner and Elser, 2002; Lavrinienko et al., 2021), and DNA contents correlate with mycelial growth rates in isolates tested her (unpublished data). As a consequence, environmental factors selecting for certain fungal strategies may also affect ITS copies present in soil fungal communities.

### Open questions in quantifying fungi in soil

Beside interspecific variability and phylogenetic diversity, there remain further open questions regarding the application of conversion factors for soil fungal biomarkers: (1) the comparability of fungal marker contents in laboratory analyses compared to fungi living in their natural soil habitat; (2) age-related shifts in biomarker contents and its physiological significance for fungal growth stages; (3) the impact of fungal spores on soil biomarker profiles.

Relative biomarker contents in fungal biomass vary with environmental conditions (Wallander et al., 2013; Joergensen, 2022). Here, culturing media were adjusted to obtain sufficient pure fungal biomass for biomarker analyses. However, rich media based on glucose and inorganic nutrients reduce fungal expenses for enzymes (Hsieh et al., 2014), induce internal storage mechanisms (Mason-Jones et al., 2023) and likely also reduce the necessity of internal recycling of cellular components (Heaton et al., 2016). Fatty acid marker composition seems to be relatively robust to environmental changes (Stahl and Klug, 1996; Klamer and Bååth, 2004), but shifts in quantitative fungal biomarker contents have not been tested. The higher conversion factor for the fungal PLFA biomarker reported by Joergensen and Wichern (2008) based on indirect soil analyses might reflect such decreased PLFA concentration in fungi growing in soil. In the case of ergosterol, there exists more evidence for its variation with culturing conditions, nutrient and carbon supply, but results remain inconclusive (Charcosset and Chauvet, 2001; Niemenmaa et al., 2008; Song et al., 2014). While Klamer and Bååth (2004) report lower ergosterol contents on complex C media compared to simple sugars, Niemenmaa et al. (2008) found high fungal ergosterol content under C starvation. Similarly, DNA contents may be reduced under resource limitations, leading to higher conversion factors in soil (Grimmett et al., 2013; Lavrinienko et al., 2021). In conclusion, we do not only recommend determining fungal biomarker contents in more fungal species in future studies, but also when cultivated under more *soil-like* resource conditions in order to refine conversion factors even further.

Biomarker contents have been repeatedly shown to be age-related (Klamer and Bååth, 2004; Niemenmaa et al., 2008; Song et al., 2014). Mycelial growth is characterized by particular physiological mechanisms. Hyphal tip growth leads to internal recycling of cell components accompanied by vacuolization of distant/older hyphae (Camenzind et al., 2021). This physiological aging/senescence primarily affects the cytoplasm (Klein and Paschke, 2004), but also membrane and cell wall composition to a certain extent (Pusztahelyi et al., 2006). Experimental evidence does not support a consistently negative correlation of the cell membrane components PLFA and ergosterol with culturing age (Stahl and Klug, 1996; Charcosset and Chauvet, 2001; Niemenmaa et al., 2008). By contrast, DNA concentrations in fungal biomass (together with other cytoplasmic components) are reduced over time (Grimmett et al., 2013; Camenzind et al., 2021). Consequently, PLFA and ergosterol components, in contrast to DNA, may include more inactive and senescing mycelial fractions. Still, these biomarkers represent a valid estimate of fungal biomass, since turnover rates of ergosterol and PLFA are relatively high in soil (Frostegård and Bååth, 1996; Miltner et al., 2012; Ekblad et al., 2016). One conclusion may be that the combination and ratios of different fungal biomarkers provide interesting insights into fungal activity in soil: shifts in DNA (and especially RNA) : PLFA content ratios could provide insights into the inactive hyphal fraction. This can further be combined with amino sugar analyses to determine dead fungal proportions (Joergensen, 2018), or NLFA marker analyses as an indicator of fungal storage (Gorka et al., 2023).

Similar to uncertainties in sequencing studies, the contribution of fungal spores to soil biomarker contents remains unclear. Spore production - sexual but particularly asexual - is an integral part of fungal development in soil (Domsch et al., 2007). Fungal sporulation rates are high (Camenzind et al., 2022) and indirect assessments of spore contributions to colony forming units approve their abundant presence in soil (Thorn et al., 1996). The fatty acid profile in spores is comparable to hyphae, as shown for basidiospores (Brondz et al., 2004), though quantitative PLFA contents remain unknown. In arbuscular mycorrhizal fungi, the fatty acid profile (with 16:1ω5 as marker for arbuscular mycorrhizal fungi) is similar to hyphae (Olsson and Johansen, 2000), but spores were enriched in storage NLFA. The PLFA marker 16:1ω5 is therefore frequently used to address the occurrence of growing hyphae in comparison to spores (Ngosong et al., 2012). Independent of exact values, fungal spores do contain relevant concentrations of fungal biomarkers, including ergosterol and DNA (Pasanen et al., 1999). As a consequence, soil fungal biomarkers not only capture hyphae, but also spores present in soil, with the relative contribution depending on spore numbers present in soil.

### Conclusions and outlook

The data of our study provide an update of soil fungal biomarker conversion factors and shed light on the relevance of interspecific variability. At this stage, fungal PLFA biomarkers appear most comparable and widely applied (Yu et al., 2022). The phylogenetic coverage of PLFA 18:2ω6,9 was higher than for ergosterol, though still problematic for early fungal lineages, while interspecific variability was lower than in ITS1 copies. Potential deviations in PLFA conversion factors (and also other biomarker types) under natural soil conditions should be further evaluated to correctly identify related fungal C contents. We further highly recommend refining the application of a conversion factor for fungal qPCR data in future studies. Despite high interspecific variability - that may be negligible in naturally diverse soils - qPCR results based on ITS sequences provide the benefit of having the least phylogenetic bias and absolute specificity for fungi (in contrast to problematic biases in PLFA). Combinations of these data with stable isotope probing and sequencing data provide interesting avenues for future in depth microbial community analyses, and a better resolution for fungal guilds (i.e., saprobes, pathogens and mycorrhizae; Nuccio et al., 2022). A combined analysis of different biomarkers (including also RNA (active), NLFA (storage) and amino sugars (necromass)) would aid for a precise quantification of fungi in soil, in addition to interesting insights into the physiological status of soil fungi under different conditions (Canarini et al., 2023).

To conclude, the precise quantification of microbial groups is essential for understanding soil C dynamics (He et al., 2021). Biomarker analyses provide the best proxy for fungal and bacterial biomass, still, an improved understanding is needed to fully exploit the potential of these data. Our data add relevant aspects on the variability in biomarker contents among fungal communities, and improve the conversion to fungal C contents. Such methodological insights are needed to improve the mechanistic understanding of underlying microbial physiology, which should be optimized prior to their inclusion in global soil C models.

## Acknowledgments

TC acknowledges funding by the Deutsche Forschungsgemeinschaft (DFG, grant number 465123751, SPP2322 SoilSystems). SH was partly supported by DFG grant HE 6183/5-1 and SM by MA4436/1-5. We thank Alberto Canarini and Kyle Mason-Jones for important insights on fungal storage mechanisms.

