## Supporting Information S1 for "Revisiting soil fungal biomarkers and conversion factors: Interspecific variability in phospholipid fatty acids, ergosterol and rDNA copy numbers"

Journal: submitted to *Soil Ecology Letters*

Authors: Camenzind, T, Haslwimmer, H, Rillig, MC, Ruess, L, Finn, D[5], Tebbe, C, Hempel, S, Marhan, S

**Table S1** Details of the analyzed fungal isolates including taxonomic classification based on long sequence reads of ITS1, 5.8S, ITS2 and partial LSU regions (including DSMZ accession numbers (German Collection of Microorganisms and Cell Cultures GmbH); phylogeny according to Unite database (Nilsson et al., 2018))

| **strain ID** | **DSMZ accession number** | **phylum** | **class** | **order** | **family** | **taxon name** |
|---|---|---|---|---|---|---|
| RLCS01 | DSM100293 | Mucoromycota | Mucoromycetes | Mucorales | Mucoraceae | *Mucor fragilis* |
| RLCS02 | DSM100407 | Mortierellomycota | Mortierellomycetes | Mortierellales | Mortierellaceae | *Mortierella sp.3* |
| RLCS03 | DSM100285 | Mortierellomycota | Mortierellomycetes | Mortierellales | Mortierellaceae | *Mortierella alpina2* |
| RLCS04 | DSM100322- | Mortierellomycota | Mortierellomycetes | Mortierellales | Mortierellaceae | *Mortierella sp.2* |
| RLCS05 | DSM100403 | Ascomycota | Sordariomycetes | Hypocreales | Nectriaceae | *Fusarium sp.1* |
| RLCS06 | DSM100400 | Ascomycota | Sordariomycetes | Sordariales | Chaetomiaceae | *Chaetomium angustispirale1* |
| RLCS07 | DSM100284 | Ascomycota | Sordariomycetes | Xylariales | Bartaliniaceae | *Truncatella angustata* |
| RLCS08 | DSM100325 | Ascomycota | Sordariomycetes | Hypocreales | Nectriaceae | *Fusarium sp.3* |
| RLCS09 | DSM100406 | Basidiomycota | Agaricomycetes | Polyporales | Polyporaceae | *Trametes versicolor* |
| RLCS10 | DSM100286 | Ascomycota | Dothideomycetes | Pleosporales | Pleosporaceae | *Alternaria sp.* |
| RLCS11 | DSM100289 | Mortierellomycota | Mortierellomycetes | Mortierellales | Mortierellaceae | *Mortierella alpina1* |
| RLCS12 | DSM100405 | Ascomycota | Sordariomycetes | Sordariales | Chaetomiaceae | *Chaetomium angustispirale2* |
| RLCS13 | DSM100290 | Ascomycota | Sordariomycetes | Hypocreales | Nectriaceae | *Fusarium sp.2* |
| RLCS14 | DSM100404 | Ascomycota | Dothideomycetes | Pleosporales | Didymellaceae | *Nothophoma sp.* |
| RLCS15 | DSM100402 | Mortierellomycota | Mortierellomycetes | Mortierellales | Mortierellaceae | *Mortierella sp.1* |

| | | | | | | |
|---|---|---|---|---|---|---|
| RLCS16 | DSM100408 | Basidiomycota | Agaricomycetes | Agaricales | Pleurotaceae | *Pleurotus sp.* |
| RLCS17 | DSM100324 | Basidiomycota | Agaricomycetes | Agaricales | Entolomataceae | *Clitopilus sp.* |
| RLCS18 | DSM100287 | Ascomycota | Sordariomycetes | Hypocreales | Nectriaceae | *Fusarium gibbosum* |
| RLCS19 | DSM100331 | Mucoromycota | Umbelopsidomycetes | Umbelopsidales | Umbelopsidaceae | *Umbelopsis isabellina* |
| RLCS20 | DSM100329 | Ascomycota | Sordariomycetes | Hypocreales | Ophiocordycipitaceae | *Purpureocillium lilacinum* |
| RLCS21 | DSM100327 | Ascomycota | Dothideomycetes | Pleosporales | Cucurbitariaceae | *Pyrenochaetopsis leptospora* |
| RLCS22 | DSM100401 | Ascomycota | Dothideomycetes | Pleosporales | Phaeosphaeriaceae | *Paraphoma chrysanthemicola* |
| RLCS23 | DSM101519 | Ascomycota | Sordariomycetes | Hypocreales | Stachybotryaceae | *Paramyrothecium sp.* |
| RLCS24 | DSM100410 | Ascomycota | Sordariomycetes | Hypocreales | Clavicipitaceae | *Metarhizium marquandii* |
| RLCS25 | DSM100292 | Ascomycota | Sordariomycetes | Hypocreales | Bionectriaceae | *Gliomastix sp.* |
| RLCS26 | DSM100330 | Ascomycota | Leotiomycetes | Helotiales | Helotiaceae | *Tetracladium apiense* |
| RLCS27 | DSM100326 | Ascomycota | Sordariomycetes | Sordariales | Chaetomiaceae | *Chaetomium subspirilliferum* |
| RLCS28 | DSM100323 | Ascomycota | Leotiomycetes | Helotiales | NA | *NA* |
| RLCS30 | DSM100291 | Ascomycota | Eurotiomycetes | Chaetothyriales | Herpotrichiellaceae | *Exophiala sp.* |
| RLCS31 | DSM100328 | Ascomycota | Eurotiomycetes | Chaetothyriales | Cyphellophoraceae | *Cyphellophora sp.* |

Soil fungal community analyses in selected soils used as a template for modeling

Three soils were chosen used as standard soils in the priority program SoilSystems (https://soilsystems.net/about/common-experimental-platform/soils/), originating from long-term field experiments. Fertilized and unfertilized soils from the Dikopshof site (University Bonn, Germany; referred to as *Soil1* and *Soil2*), and fertilized soil from Thyrow (Humboldt-Universität Berlin, Germany; *Soil3*) were chosen. Soils were stored at -20° until analyses. Five hundred milligrams of soil was used for DNA extraction via the FastDNA Spin Kit (MP Biomedicals, Eschwege, Germany) as per the manufacturer's recommendations. ITS copy numbers were determined by qPCR on 10 ng $\mu L^{-1}$ standardised DNA template, using primers NSI1 and 58A2R (Martin and Rygiewicz, 2005) in SYBR green reactions (Thermo Fisher, Osterode am Harz, Germany) on a CFX96 Touch realtime PCR detection system (Biorad laboratories, Neuried, Germany) as described previously (Hemkemeyer et al., 2015) Standard curves were derived from known quantities of *Fusarium culmorum*, and qPCR efficiency was 93.5%, $R^2$ of 98.3%.

For fungal community analyses, partial fungal ITS1 sequences were amplified with barcoded ITS1f and ITS2 primers (Walters et al., 2015; Hoggard et al., 2018) in 25 µL PCR reactions using Q5 high-fidelity DNA polymerase (New England Biolabs, Ipswich, MA, USA) as described previously (Finn et al., 2023). Amplicon products were purified and normalized to 2 ng $\mu L^{-1}$ prior to library prep with Ovation Rapid DR Multiplex System 1-96 (NuGEN Technologies, San Carlos, CA, USA) and sequencing on the Illumina MiSeq platform by LGC Genomics GmbH, Berlin, Germany). Sequences were analyzed using the DADA2 ITS Pipeline Workflow (1.8) (https://benjjneb.github.io/dada2/ITS_workflow.html; Callahan et al., 2016). Sequences were assigned to phyla using the UNITE database (Nilsson et al., 2018).

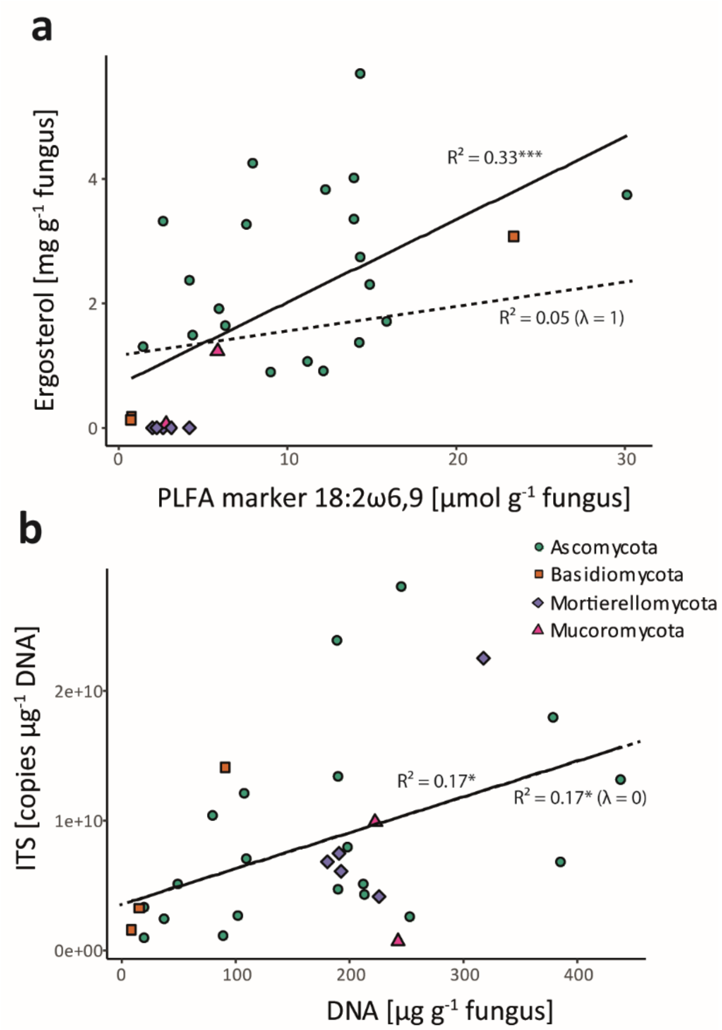


**Fig. S1** Significant linear correlations of fungal biomarker contents in individual fungal isolates. Solid lines represent linear regression models, dashed lines phylogenetically corrected linear correlations (phylogenetic generalized least squares (PGLS)). Correlation coefficients ($R^2$), respective P-values (* $P < 0.05$, *** $P < 0.001$) and phylogenetic signals are given (pagel´s lambda λ). Symbol shape and colour reflect respective phyla affiliations (see legend).
